## Supplementary Figures 1 through 9 for "DNA binding analysis of rare variants in homeodomains reveals novel homeodomain specificity-determining residues"

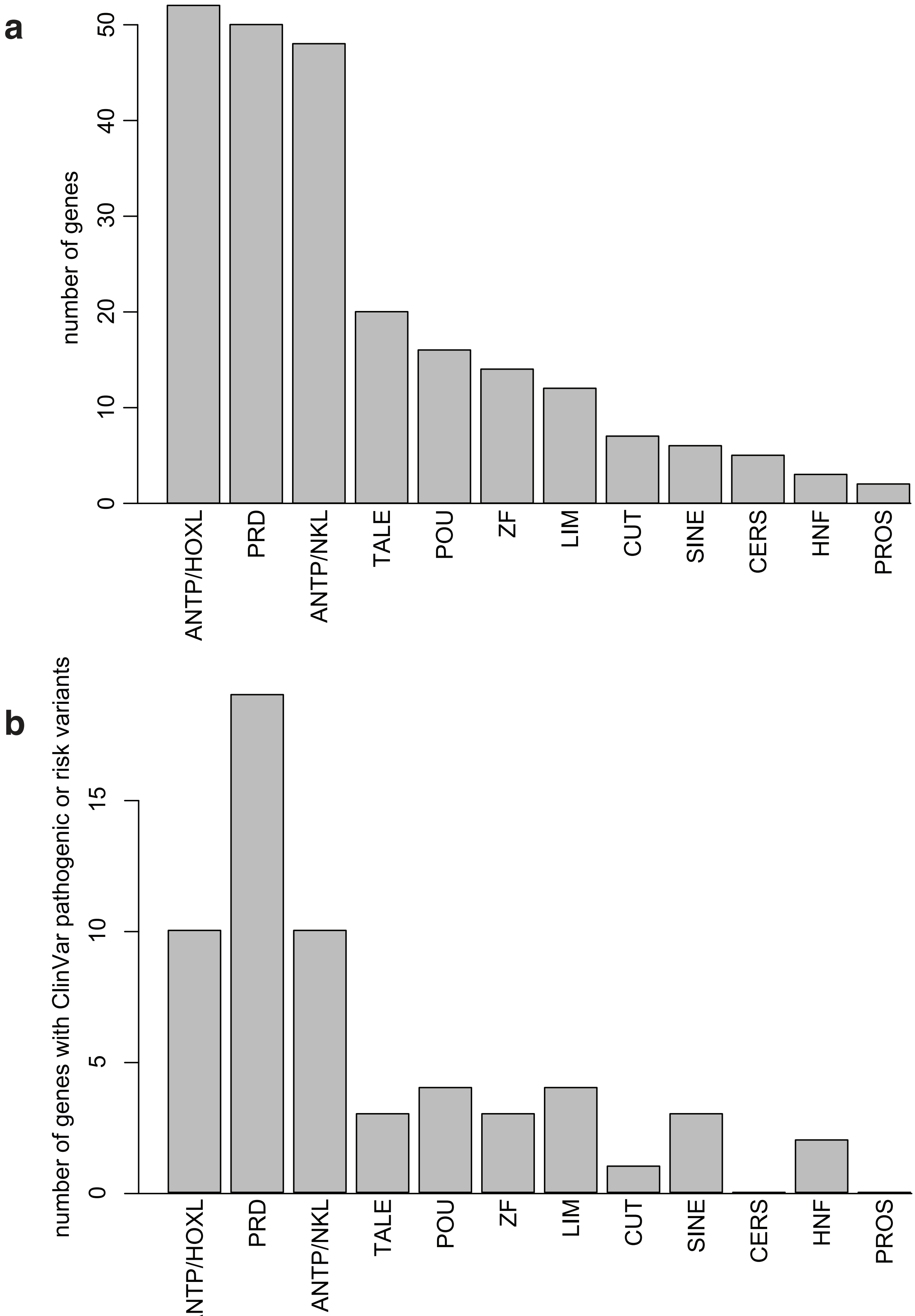


**Supplementary Fig. 1: (a)** Number of genes in each HD subfamily. **(b)** Number of genes with ClinVar disease alleles (pathogenic, likely pathogenic, and/or risk) in each HD subfamily.

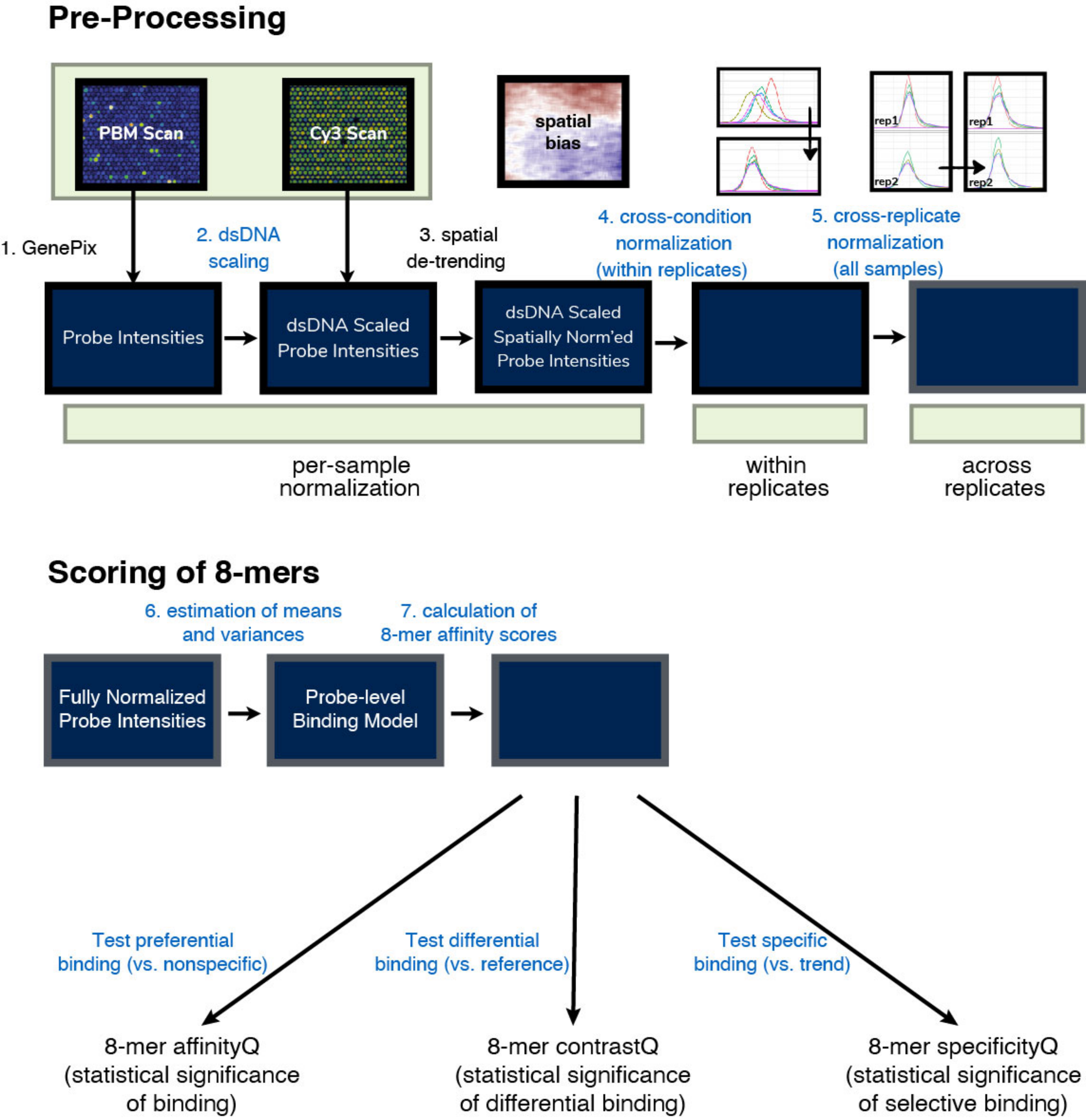


**Supplementary Fig. 2:** Schema of uPBM analysis. Steps indicated in black are part of the previously published workflow [70,71]; steps indicated in blue are new in this study.

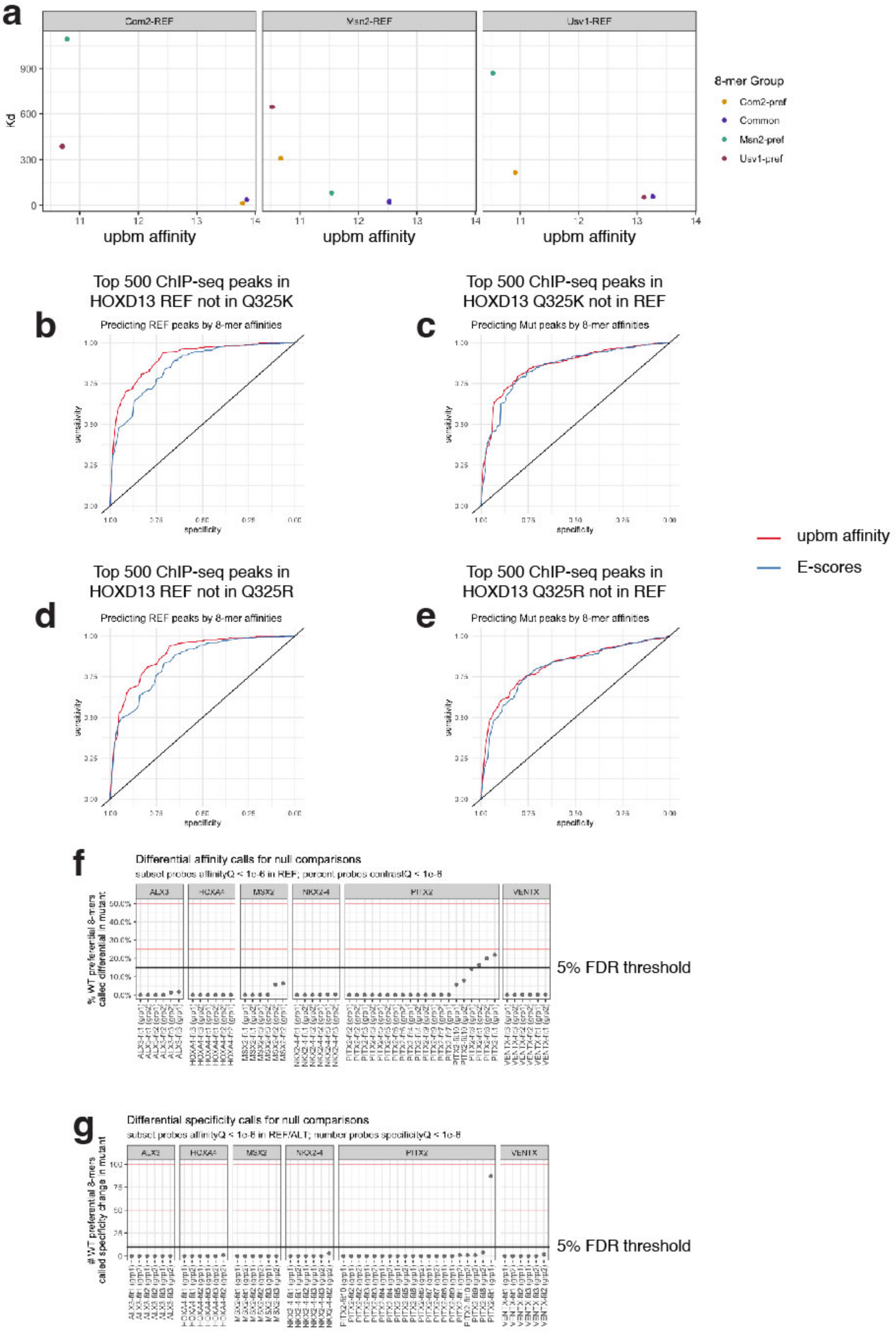


**Supplementary Fig. 3**: Validations of upbm analysis method. **(a)** Siggers Mol Cell 2014 paper [24]. **(b-e)** HOXD13 mutants [17]. **(f)** Fraction of significantly bound 8-mers with significantly altered affinity in null (reference vs. reference) comparisons and selection of 5% FDR threshold for calling affinity-changing variants. **(g)** Number of significantly bound 8-mers with significant specificity in null (reference vs. reference) comparisons and selection of 5% FDR threshold for calling specificity-changing variants.

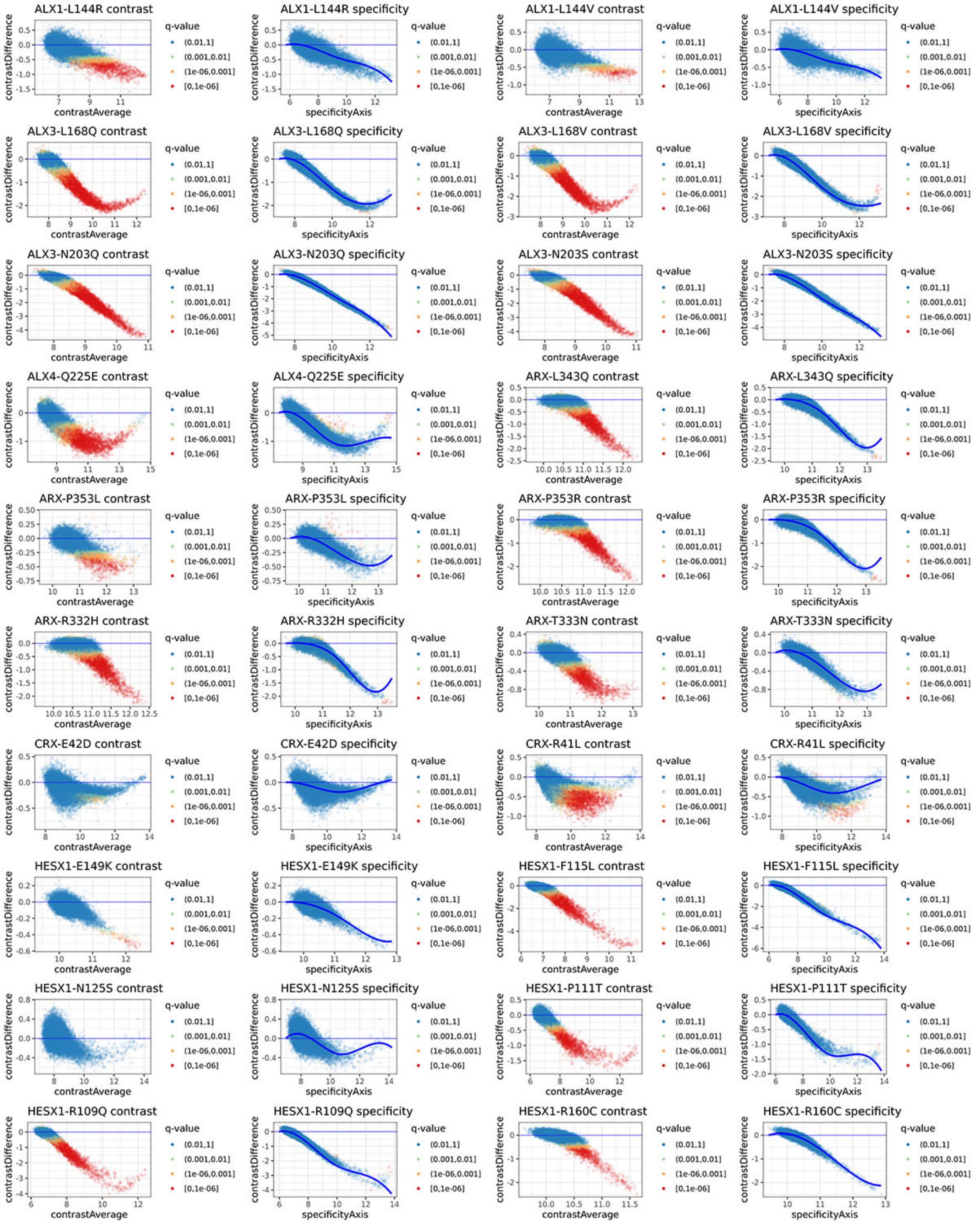


**Supplementary Fig. 4:** Differential 8-mer MA plots colored by contrastQ (left) and specificityQ (right) for all variants.

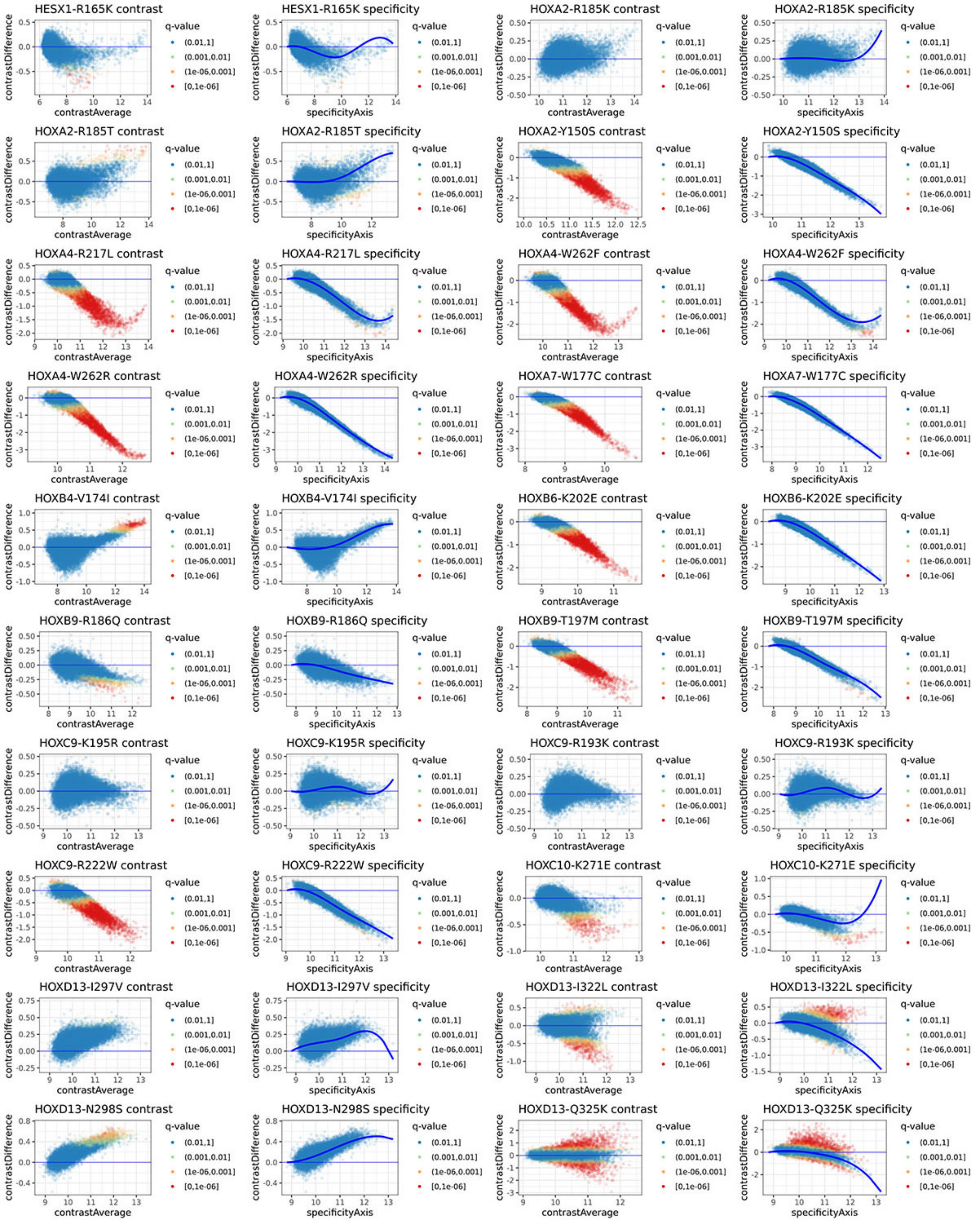


**Supplementary Fig. 4**, continued.

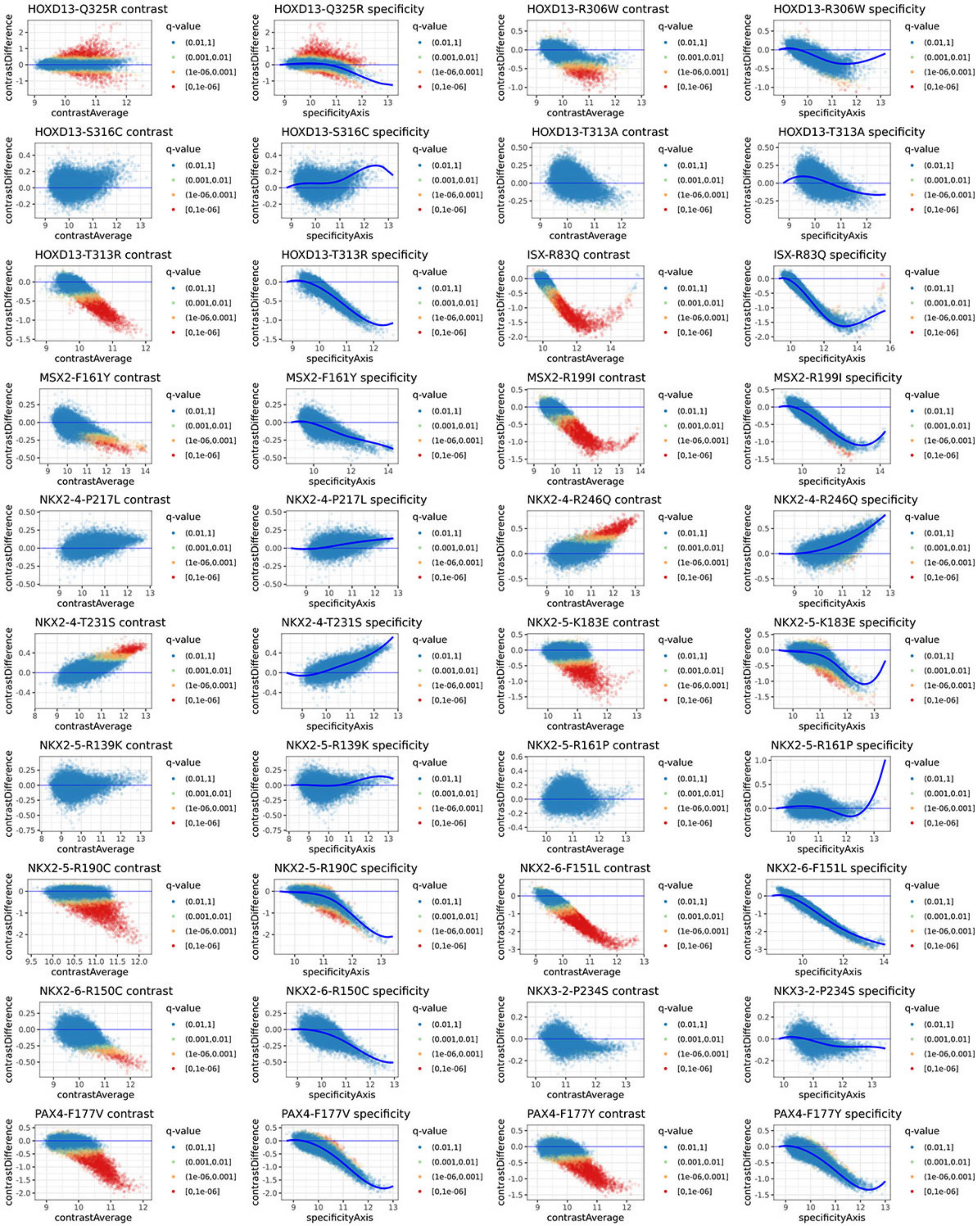


**Supplementary Fig. 4**, continued.

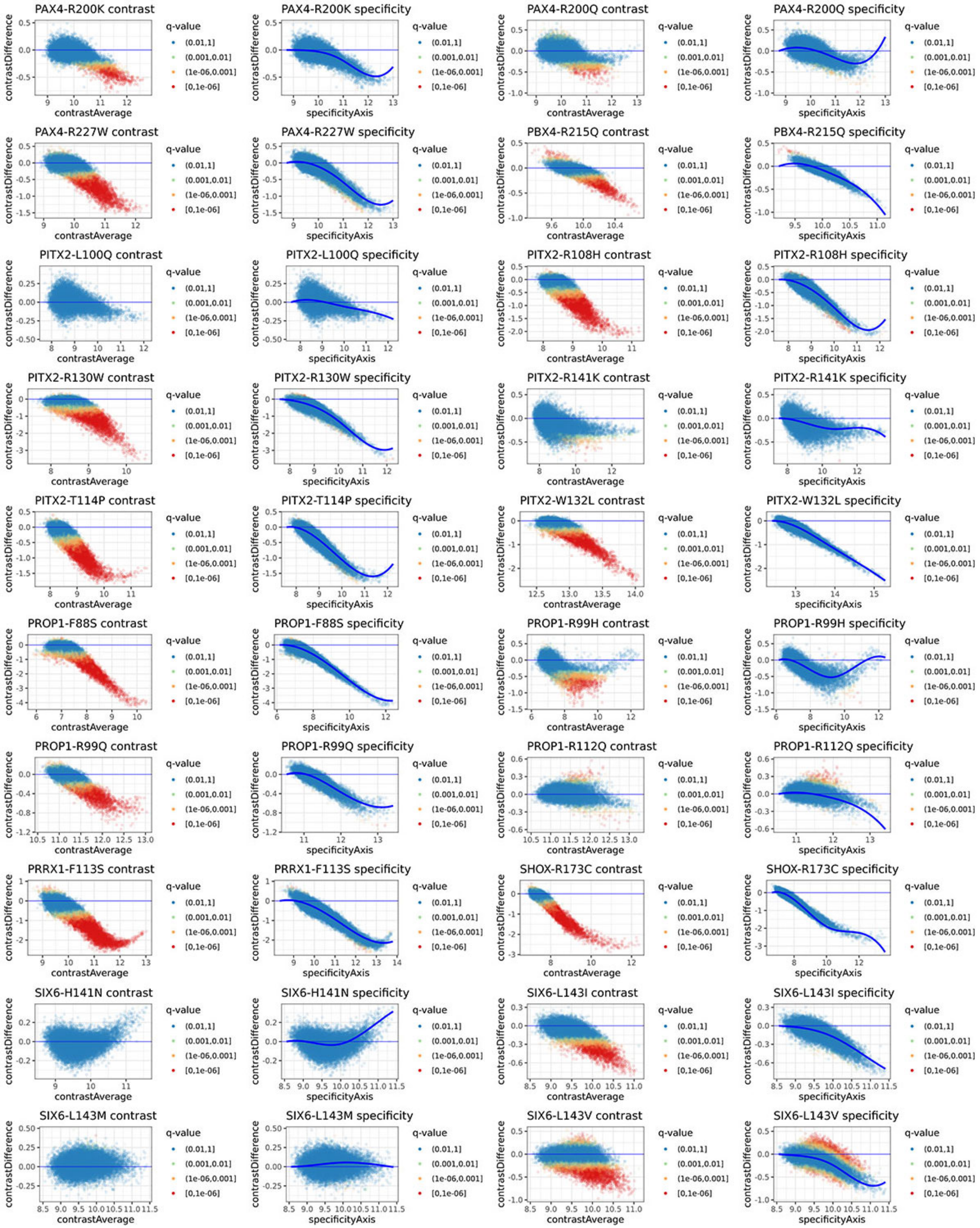


**Supplementary Fig. 4**, continued.

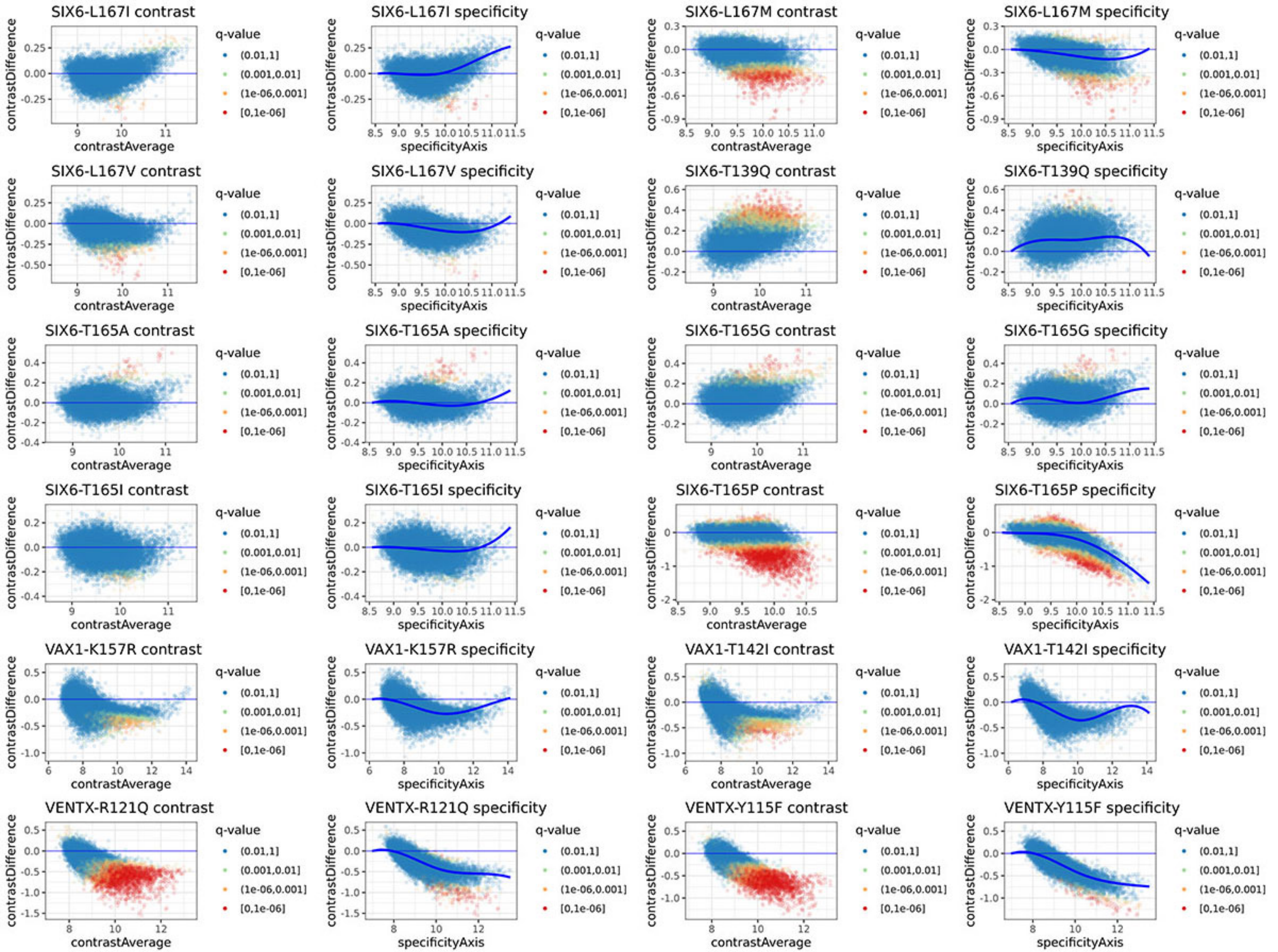


**Supplementary Fig. 4**, continued.

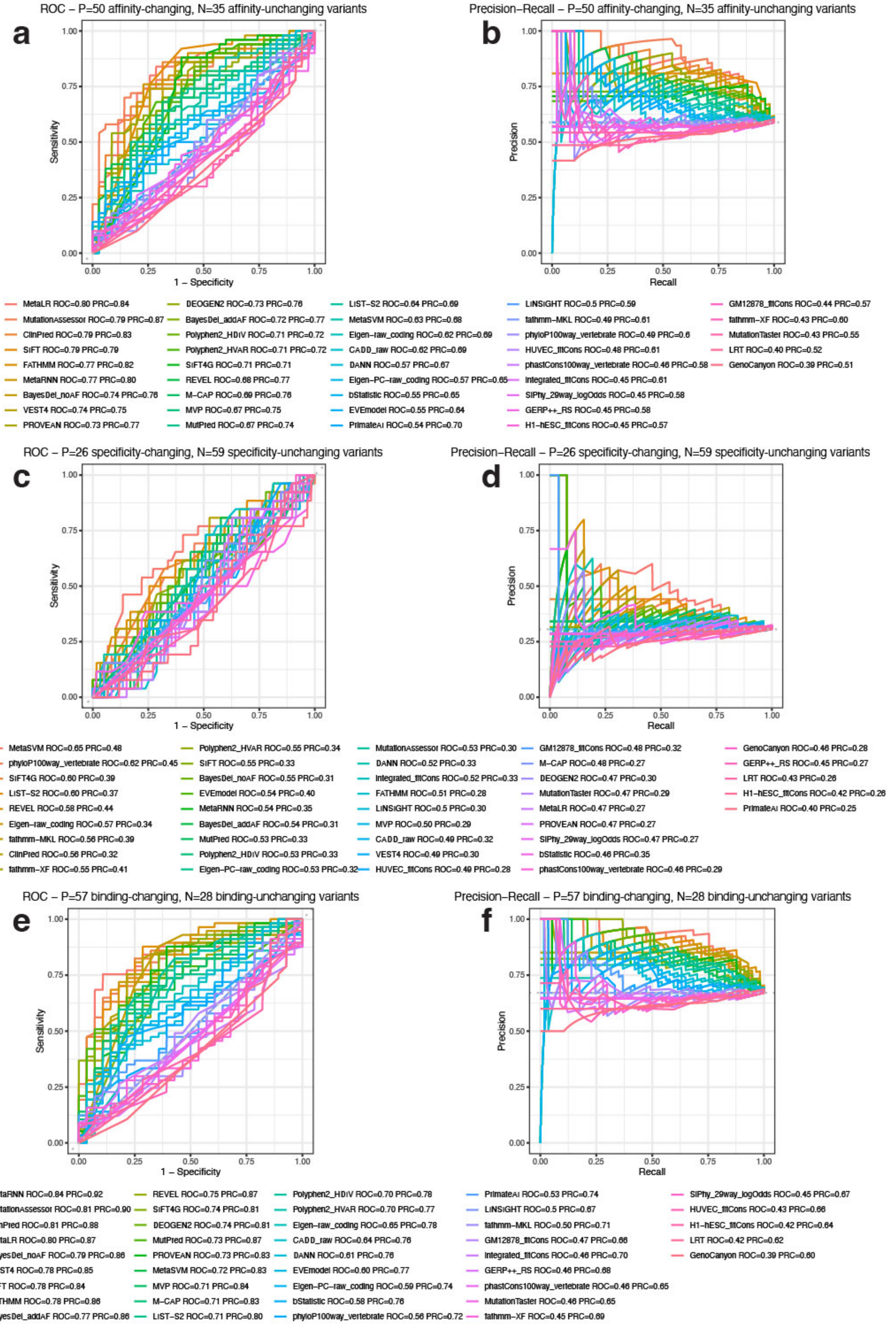


**Supplementary Fig. 5:** AUROC and AUPRC plots for all variant effect prediction tools for discriminating variants with altered **(a,b)** affinity, **(c,d)** specificity, or **(e,f)** binding (*i.e.,* affinity and/or specificity).

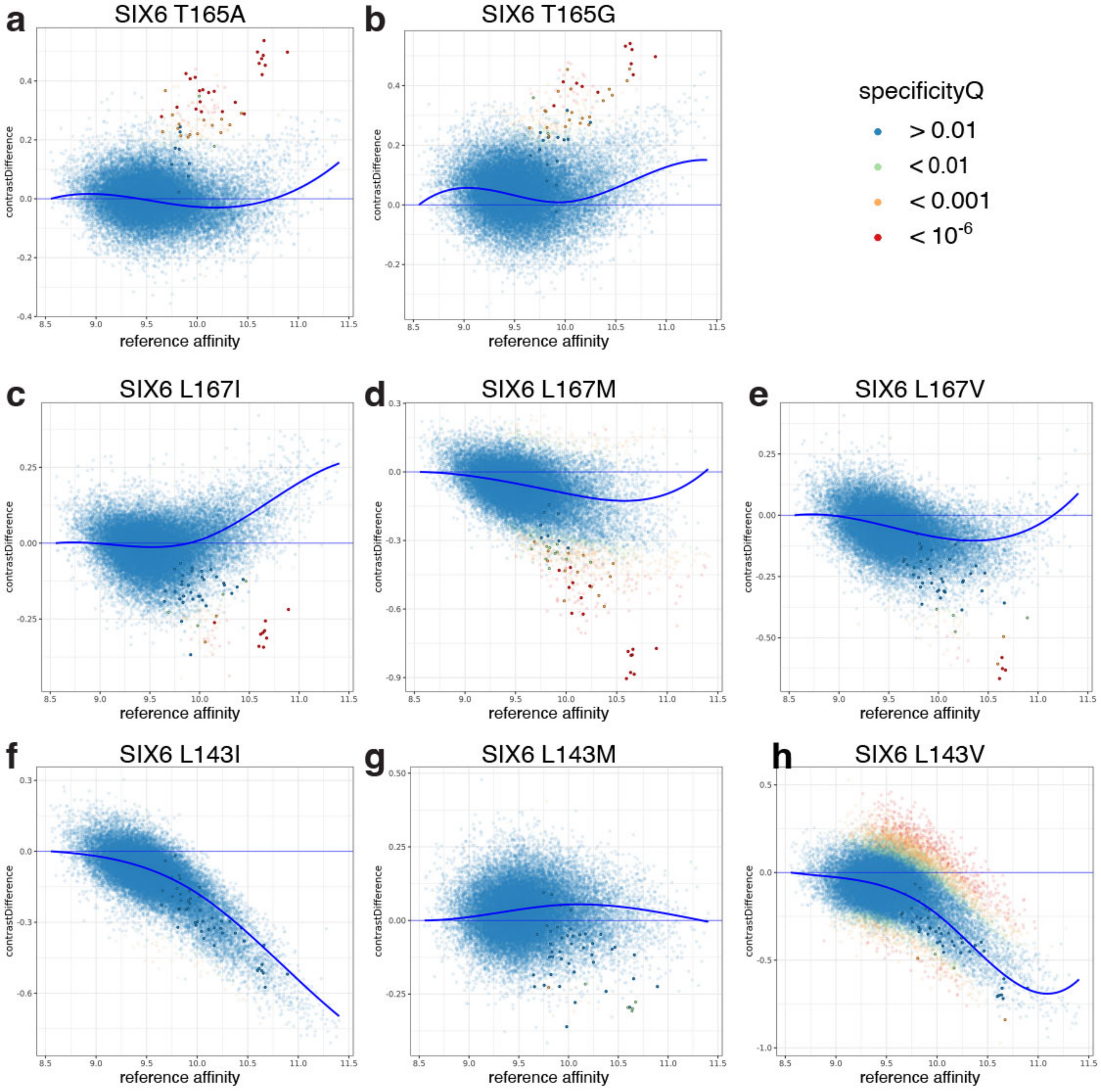


**Supplementary Fig. 6:** Amino acid positions distal to the DNA-binding interface affect specificity in SIX6. The **(a)** T165A and **(b)** T165G variants preferentially increase affinity for 8-mers containing a TGACAC motif (highlighted points). The **(c)** L167I, **(d)** L167M, and **(e)** L167V variants preferentially decrease binding to the same 8-mers. The **(f)** L143I, **(g)** L143M, and **(h)** L143V variants have more variable effects.

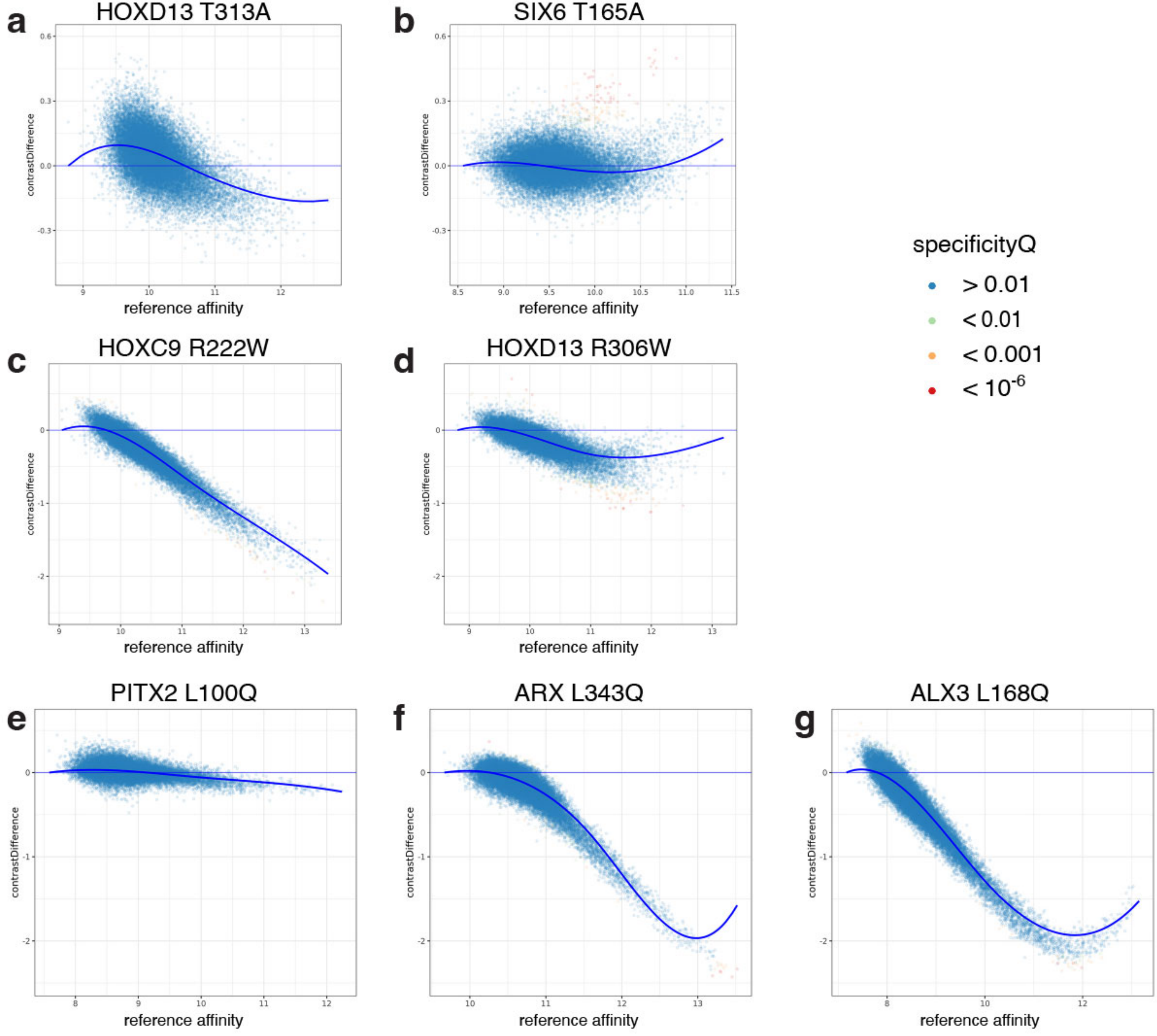


**Supplementary Fig. 7:** Same substitution in different HDs showed different effects on DNA binding activity. **(a)** Thr to Ala substitution at HD canonical position 38 resulted in no change in DNA binding activity in HOXD13, but **(b)** altered specificity in SIX6. **(c)** Arg to Trp substitution at canonical position 31 resulted in strongly reduced affinity in HOXC9, an ANTP/HOXL HD, but **(d)** mildly reduced affinity and altered specificity in HOXD13, another ANTP/HOXL HD. **(e)** Leu to Gln substitution at position 16 showed no effect in PITX2, but resulted in strongly decreased affinity in **(f)** ARX and **(g)** ALX3.

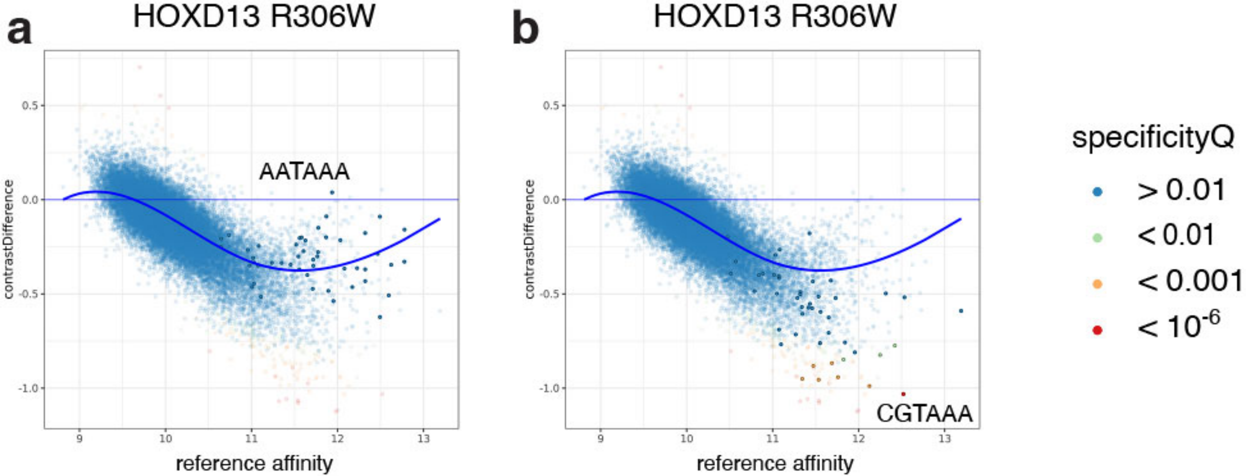


**Supplementary Fig. 8:** HOXD13 R306W variant appears to differentially affect binding to **(a)** AATAAA- vs. **(b)** CGTAAA-containing 8mers.

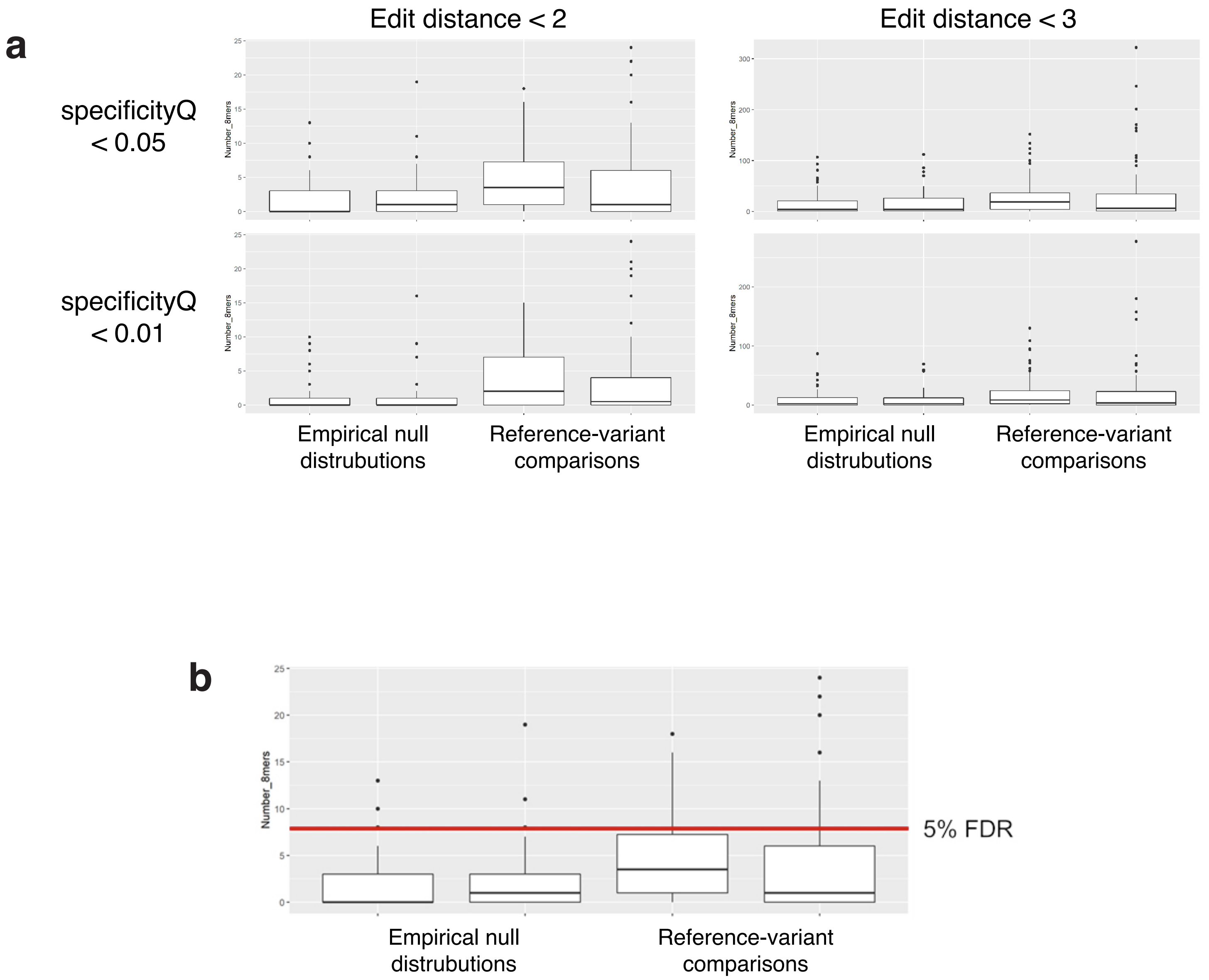


**Supplementary Fig. 9: (a)** Distribution of numbers of 8-mers with specificityQ below threshold within an edit distance below threshold of the top differentially bound 8-mer, for various thresholds, in null (reference vs. reference) comparisons (left) or variant vs. reference comparisons (right). **(b)** Example of the choice of a 5% FDR threshold for calling specificity-altering variants, in this case for specificityQ < 0.05 and edit distance < 2.
